# PatternJ: an ImageJ toolset for the automated and quantitative analysis of regular spatial patterns found in sarcomeres, axons, somites, and more

**DOI:** 10.1101/2024.01.17.576053

**Authors:** Mélina Baheux Blin, Vincent Loreau, Frank Schnorrer, Pierre Mangeol

## Abstract

Regular spatial patterns are ubiquitous forms of organization in nature. In animals, regular patterns can be found from the cellular scale to the tissue scale, and from early stages of development to adulthood. To understand the formation of these patterns, how they form and mature, and how they are affected by perturbations, a precise quantitative description of the patterns is essential. However, accessible tools that offer in-depth analysis without the need for computational skills are lacking for biologists. Here we present PatternJ, a novel toolset to analyze regular pattern organizations precisely and automatically. This toolset, to be used with the popular imaging processing program ImageJ/Fiji, facilitates the extraction of key geometric features within and between pattern repeats. We validated PatternJ on simulated data and tested it on images of sarcomeres in insect muscles and cardiomyocytes, actin rings in neurons, and somites in zebrafish embryos obtained using confocal fluorescence microscopy, STORM, electron microscopy, and bright-field imaging. We show that the toolset delivers subpixel feature extraction reliably even with images of low signal-to-noise ratio. PatternJ’s straightforward use and functionalities make it valuable for various scientific fields requiring quantitative pattern analysis, including the sarcomere biology of muscles or the patterning of mammalian axons, speeding up discoveries with the bonus of high reproducibility.

## Main text

Regular patterns are a very common type of organization in animals. The division of the animal body plan into regular body segments is crucial for animal development from flies to humans (Hubaud & Pourquié, 2014; Nüsslein-Volhard & Wieschaus, 1980). Within each segment, bristles organize regularly at the surface of adult insects (Lawrence et al., 1979; Orenic et al., 1993), such as feathers do on the skin of birds (Haupaix et al., 2018), or scales on the skin of reptiles (Chang et al., 2009). At the cellular scale, regularly repeated actin rings are found in mature axons (Leterrier et al., 2015; Xu et al., 2013), and actin organizes with myosin into periodic repeats in cell lines grown on rigid surfaces *in vitro* (Hu et al., 2017). Most impressively in muscles, proteins organize in periodic sarcomeres that repeat themselves very regularly up to a hundred thousand times along a single myofibril (Lange et al., 2006; Luis & Schnorrer, 2021).

To understand how these regularly repeated patterns emerge over time, and how they are affected by mutations or environmental changes, it is important to precisely extract various important geometrical features, such as the size of a single pattern, its shape, or other geometrical arrangements at the level of a single pattern. Such quantitative extraction from images is often wanted by biologists. However, reaching this goal is often prone to error and bias when done manually, and it can be very time-consuming for biologists without extensive experience in computer programming.

Several automated tools have been developed in recent years to respond to such a need. The muscle field has been particularly active, with about 10 tools published over the last decade (Czirok et al., 2017; Lee et al., 2015; Morris et al., 2020; Neininger-Castro et al., 2023; Pasqualin et al., 2016; Sala et al., 2018; Spletter et al., 2018; Toepfer et al., 2019; Zhao et al., 2021). These tools are centered on the use in muscles and their sarcomeres: some focus on muscle contraction and give as output the contraction state of the tissue (Czirok et al., 2017; Lee et al., 2015; Sala et al., 2018), some can segment single sarcomeres from an image and extract their sarcomere lengths (Neininger-Castro et al., 2023; Toepfer et al., 2019) while others extract a global sarcomere length from an image with little to no input from the user (Morris et al., 2020; Pasqualin et al., 2016; Spletter et al., 2018). Often the input image is expected to display the periodic repetition of isolated single bands, usually obtained by staining of the sarcomeric Z-disk, a thin protein-rich region bordering neighboring sarcomeres. The simplest tools usually require limited to no coding knowledge (Pasqualin et al., 2016; Sala et al., 2018; Spletter et al., 2018), while the most developed ones usually require advanced programming knowledge to be implemented by the user (Czirok et al., 2017; Lee et al., 2015; Morris et al., 2020; Neininger-Castro et al., 2023; Toepfer et al., 2019; Zhao et al., 2021).

Currently, only sarcApp provides an in-depth analysis of pattern features (Neininger-Castro et al., 2023). While the other ones can provide at best the spatial periods of individual patterns, sarcApp can extract the position of staining edges within patterns with, to our knowledge, pixel precision, and offers several other analyses including the direction of patterns, the number of sarcomeres or myofibrils in an image and more. However, an important limit is that despite the efforts to make it user-friendly, it still requires programming skills that are not compatible with most users’ knowledge, and as it is based on deep learning, it will require a computer that is adapted to the task, including a powerful GPU. Hence, the field would greatly benefit from a tool that provides an in-depth analysis of pattern features, with subpixel precision, necessary to analyze fine pattern perturbation, and that works directly, without any programming knowledge, neither requiring a dedicated powerful computer to run.

In this manuscript, we present PatternJ, a novel toolset to extract geometrical features from images of regular patterns. This macro toolset for ImageJ/Fiji (Schindelin et al., 2012; Schneider et al., 2012) does not require any coding knowledge and is aimed to be particularly user-friendly. It can extract with subpixel precision the size of individual periodic pattern repeats. Within a pattern, it can extract the position of multiple bands, blocks, and the shape characteristics of a typical phalloidin actin staining in muscle, a very common structure in the muscle field that is important to quantify. The extraction of these simple geometrical features will address most quantitative needs when analyzing repeated patterns found in nature. Finally, the tool provides the image of an averaged pattern, which is very useful for visualizations, especially for low signal-to-noise ratio (SNR) applications, as averaging will reduce noise and potentially display features that are not possible to observe otherwise. We tested the tool for robustness against low SNR, aperiodicity, and intensity fluctuation, and evaluated how it behaves with different user selections. We then tested it on images of repeated patterns obtained by us from muscles and by colleagues from various tissues with different imaging methods.

### PatternJ workflow

After a straightforward installation by the user, the toolset functions are accessible on the ImageJ/Fiji graphical user interface (Fig. 1A). The functions guide the user to analyze in a few steps the repeated patterns present in the selected image. The typical steps the user will follow are (Fig. 1):

1. **Manual selection**: Draw a selection as a line or a curve on the image and check the corresponding intensity profile (“Check your intensity profile(s)” function). Multicolor images will give a graph with one profile per color.
2. Visually check the profile.
3. **Setting pattern features**: Set the pattern characteristics through a popup window (“Set parameters of your analysis” function). The possible pattern characteristics are chosen between predefined patterns such as individual bands, blocks, or sarcomeric actin. The choice of pattern is asked only once. Note: strictly, a pattern is any region in the profile that repeats itself when translated by one pattern length, which can make the interpretation unnecessarily complex. Here the algorithm defines one pattern by having the features of highest intensity in its center, or the center of multiple bands such as in the example of Fig 1B (the configuration in which bands are the closest is used).
4. **Automated feature extraction**: The algorithm automatically finds the pattern features based on the previous settings step and the user checks visually if they are found as expected (“Visualize the extracted position” function).
5. Once the features are correctly extracted, the user can save them (“Extract and Save” function). At this stage, steps 1, 2, 4, and 5 can be repeated on other selections in the image, or in other images.
6. **Analysis**: After repeating these steps for multiple selections on one or multiple images, the user can then proceed to the analysis of the patterns (“Analysis” function). The function will compute the pattern period for each pattern individually, save it in a file, and concatenate the features extracted in separate files for each channel. Finally, it will compute from all patterns analyzed the average intensity profile of a pattern and an image of the average pattern.

**Figure 1:**
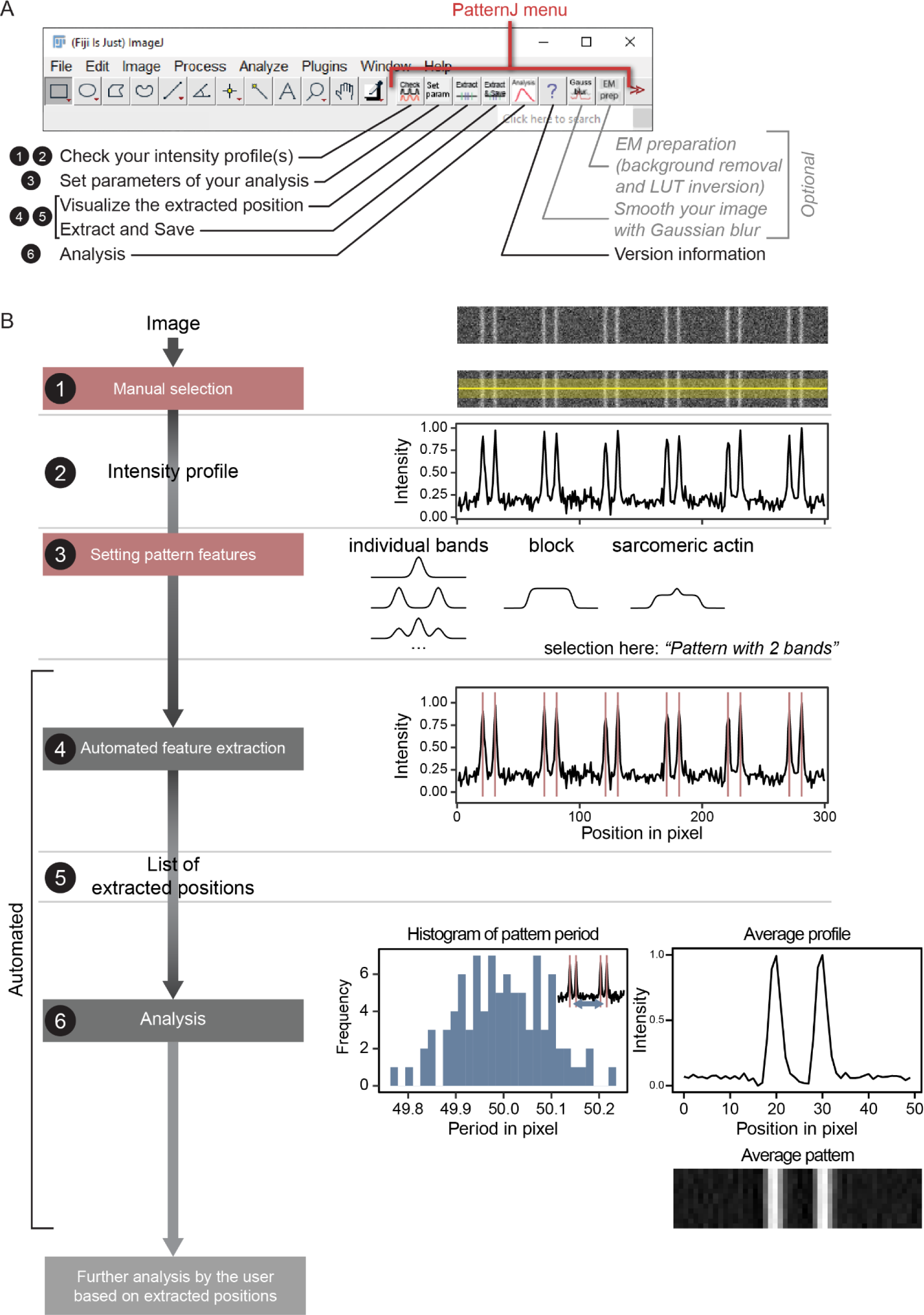
PatternJ main features. (**A**) Graphical user interface of PatternJ in Fiji and list of functions. The numbers 1 to 6 are the ones of the steps presented in (B). (**B**) Analysis steps followed by the user with the help of functions listed in (A). Detailed description in the text.

The user can further analyze the positions of the pattern features extracted as they can be found in user-friendly generated text files.

In addition to these steps, we added two optional steps. The first one uses a Gaussian blur on the image in case the image presents a very low SNR; we found it to be useful, although it does lower the image resolution slightly. The second option inverts the gray values of the image. This is particularly useful when working with images from electron microscopy, which present dark values in electron-dense regions. In fluorescence imaging, this will also be useful when the feature of interest is dark in comparison to the rest of the selection.

Automated feature extraction is the core of the tool. The algorithm takes multiple steps to achieve this (Fig. S1):

- **Step 1**: The intensity profile is used to obtain a first estimate of the spatial period by identifying the first secondary peak in the autocorrelation of the intensity profile.
- **Step 2**: This estimate of the spatial period is then used to segment automatically the profile. Step 2a, the algorithm selects automatically a region that is defined as a single reference pattern: the width of the region is the spatial period defined in the previous step and its center is the point of highest intensity in the profile. Step 2b, once the reference pattern is defined, it is used to find the other patterns hidden in the profile. The algorithm computes the cross-correlation between the intensity profile of the selection and the intensity profile of the reference pattern. Step 2c, the cross-correlation profile displays multiple peaks, some of which witness the center position of patterns. To select the correct peaks, the algorithm uses the peak of highest intensity as a starting point and then searches for peak intensity values one spatial period away on each side of this starting point. Once new peaks are found, the algorithm propagates this search, looking for new peaks one spatial period away from the previous peak. Step 2d, the search results in the full segmentation of the patterns in the profile.
- **Step 3**: With the patterns segmented, the algorithm can then precisely extract features in each pattern individually. This analysis will depend on the features the user selected. If “individual band(s)” is selected, the algorithm will search for the number of bands indicated by the user and fit a Gaussian function on a band to extract precisely its center (Fig. S2). If “block(s)” is selected, a sigmoid function is used to find the edges of the block; by default, PatternJ will give as the edge the inflection point of the fit (in other words the position at which 50 % of the maximum is reached), but the parameters of the fit are given to the user who can therefore decide after the analysis on another threshold if needed. If actin is selected, which in muscle often consists of a bright central band at the Z-disk and a wider block, a combination of the two previous fitting methods is used to define the edges and the center of the actin staining.

From the user perspective, PatternJ workflow is expected to be achieved within a minute. The automated part of the algorithm should extract features with subpixel accuracy in a multiple-color image within one second or less on a typical laptop. The main input expected from the user is a manual selection on an image, which is straightforward and quick.

### Validation of PatternJ

The steps used in the algorithm are aimed at making the feature extraction robust in a large range of situations, including challenging images, in which the signal-noise-ratio (SNR) is low or which contain not very regular patterns. To test the behavior of the algorithm, we generated simulated images, for which features are known, and we varied the SNR and the position of the pattern to simulate a range as wide as possible of situations users may face.

We started by generating images with a single band pattern repeated 10 times, separated by 25 pixels (Fig. 2). All bands have the same intensity and Poisson-generated noise is added gradually to modify the SNR from 0.5 to 8. We generated 1000 images for each SNR value selected. To obtain a first estimate of the algorithm capabilities to extract the position of the bands, we used a linear selection encompassing the 10 patterns with a linewidth of 7 pixels. We evaluated the fraction of images from which all bands were correctly extracted (Fig. 2A). At SNR = 1, it is very difficult to visually locate bands and the algorithm extracted all 10 bands in only 5 % of images. However, already at SNR = 1.5, which is still very challenging visually, all 10 bands were extracted in 55 % of images. At SNR = 2, all bands were recovered in 95 % of images, and for higher SNR, all bands were recovered in all images (Fig. 2A). Next, we estimated the precision at which the algorithm extracted the position of a single band (Fig. 2A). Even in very challenging SNR conditions, the bands were found with subpixel precision. At SNR = 3, one can expect a precision of at least a quarter of a pixel, and a tenth of a pixel or better for higher SNRs. Overall, in these first examples PatternJ can extract robustly the positions of single bands even in challenging SNR conditions with a localization precision below a pixel.

**Figure 2.**
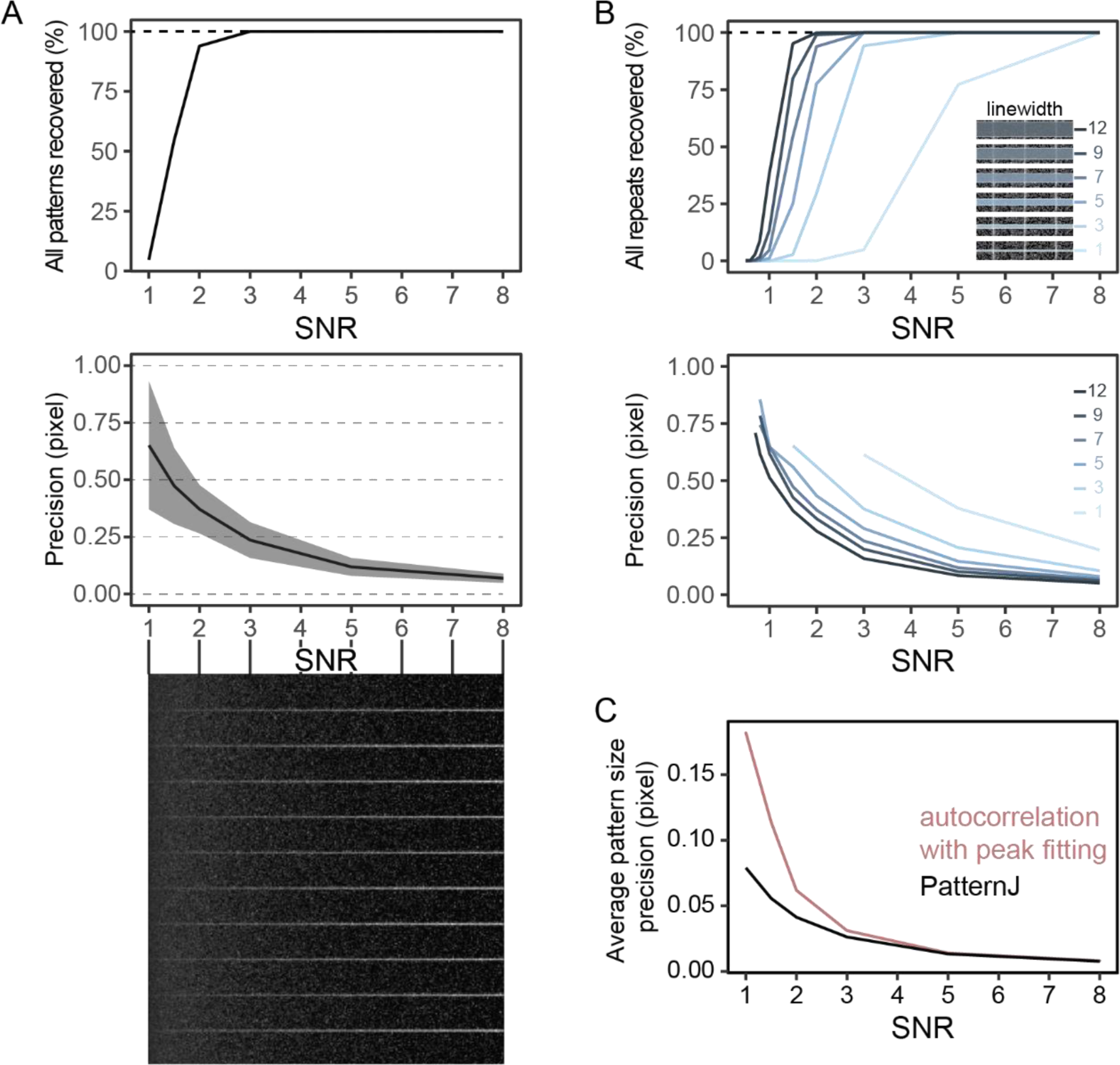
PatternJ validation on patterns with varying signal-to-noise ratio (SNR). (**A**) PatternJ performance when analyzing a single band pattern with varying (SNR). Top, fraction of selections in which all patterns were recovered. Middle, average precision in extracting the band position. Bottom, examples of pattern repeats with varying SNR. (**B**) Effect of the selection linewidth on PatternJ performance on recovery and precision. (**C**) Comparison of the precision of PatternJ and autocorrelation in extracting the pattern period. See methods for details on image generation and analysis.

Since the result of the extraction is likely to depend on the linewidth defined by the user, we changed the linewidth of the selection using the same images (Fig. 2B). The recovery and, to a lower extent, the precision were indeed affected by the linewidth chosen. With the small linewidths, the recovery and precision are reduced; however, the algorithm can still be usable in many situations a user may face. At larger linewidths, the recovery was noticeably better, even at SNR = 1. For a linewidth of 12 pixels and SNR = 1, the 10 bands were fully recovered in about 40 % of images. We also evaluated how the algorithm would behave with increasingly shorter selections. We found that the selection length has little influence on the performance of feature extraction (Fig. S3A). Altogether, we find that the algorithm robustly extracts our test pattern with high precision, even in challenging SNR conditions. Depending on the application, the user will benefit from selecting a larger linewidth, as the largest linewidths ensure the best results.

Single-band patterns are very common in images of biological samples. To analyze such patterns, it is common to manually extract the first secondary peak of the intensity profile autocorrelation or Fast Fourier Transform (FFT) to estimate the average spatial period. This analysis is usually limited to selecting the position of the peak’s highest value, which constrains the precision to one pixel. We already demonstrated that our algorithm achieves subpixel precision at the level of a single pattern. However, we were curious as to how PatternJ would compare to the autocorrelation approach if we would estimate the spatial period by precisely extracting the secondary peak of the autocorrelation function using a local Gaussian function fit; note that this improved approach is not readily accessible in Image/Fiji, but it can be achieved by writing custom programs. We found that PatternJ precision in estimating the spatial period of a pattern surpasses the improved autocorrelation approach, in particular at low SNR (Figs. 2C and S3B). The autocorrelation gives more weight to brighter bands, which biases the estimation of the mean, whereas PatternJ gives the same weight to each band, limiting bias in the mean estimation, hence the higher precision in estimating the spatial period. Therefore, we would advise using PatternJ over approaches using autocorrelation when estimating the average pattern size in which the pattern consists of a single band.

Images of biological samples of course contain more complexity than the first examples we examined. To encompass more complexity, we kept a single-band pattern and additionally varied the intensity for all bands as well as their spatial period. By varying the band intensity, we found that the band recovery was more affected by the highest level of intensity variation (Fig. 3A). We suspect that bands of lowest intensity represent the bottleneck in this analysis: at a given average SNR, these bands have a lower SNR than the other bands, which makes them more challenging to extract. When varying the spatial period, the algorithm recovered patterns well up to 20 % of variation in length, at which all 10 patterns were recovered in at least 90 % of images, especially at high SNR (Fig. 3B). At higher variation of the spatial period, where the patterns are visually less periodic, the algorithm had more difficulty in extracting patterns. As this algorithm is built to be robust for repeated patterns, this result is not surprising. However, the algorithm can still be useful in these situations. Additionally, the user must check visually the output of the algorithm, which is a default feature of PatternJ. An algorithm using a peak finder function may be better suited when patterns become aperiodic, but it is likely to require more input from the user to be reliable.

**Figure 3:**
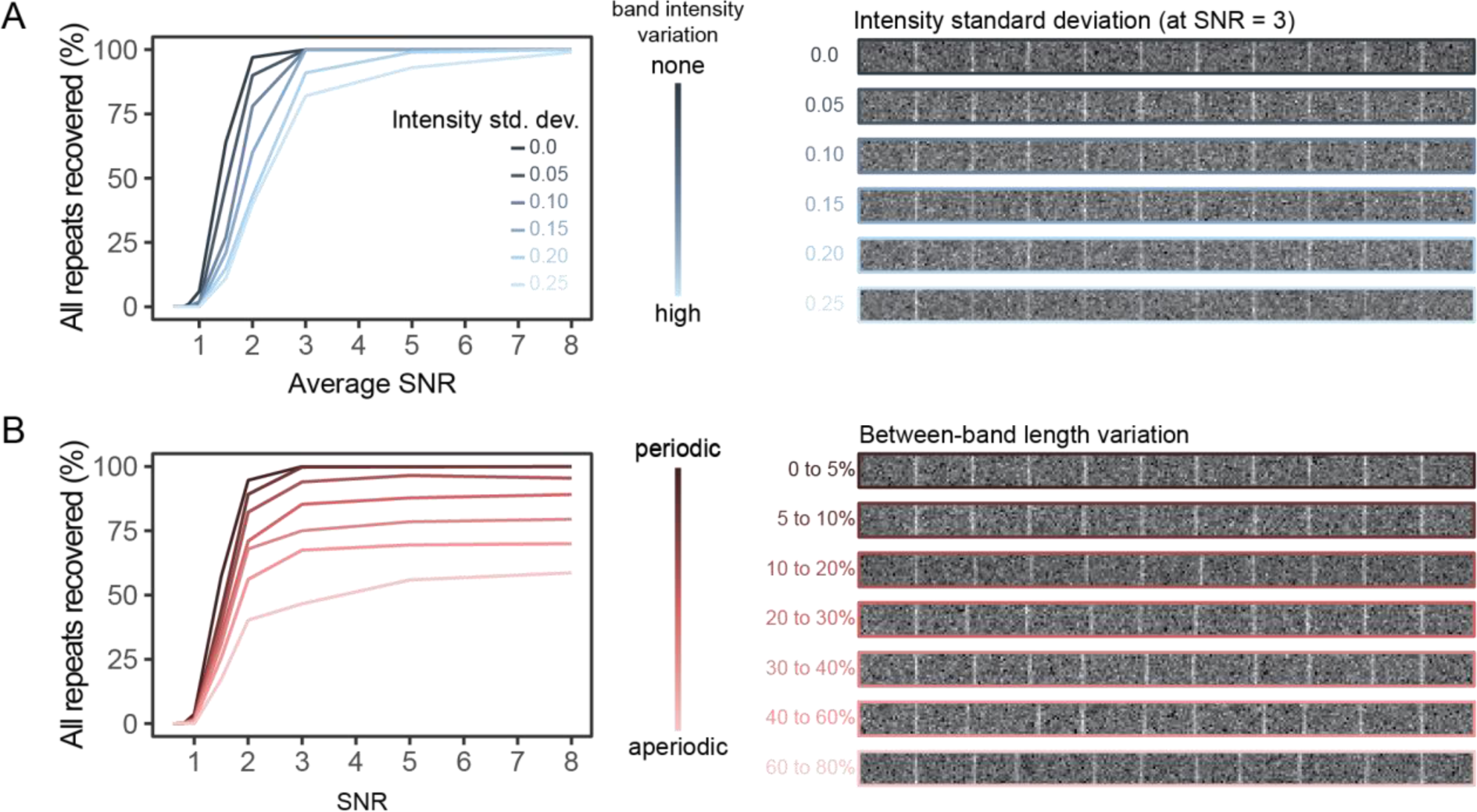
PatternJ validation on patterns with varying intensity and periodicity. (**A**). Left, effect of the band intensity variation on PatternJ performance on recovery. Right, image examples with varying band intensities. (**B**). Left, effect of the band length variation on PatternJ performance on recovery. Right, image examples with various degrees of between-band variation. See methods for details on image generation and analysis.

### Comparison of PatternJ to existing tools

Several tools to analyze repeated patterns with different aims have been published in recent years (Table 1). SarcOptiM (Pasqualin et al., 2016) and MyofibrilJ (Spletter et al., 2018) use FFT to analyze an entire selection or the entire image and compute one value for the spatial period (here sarcomere length) for the entire selection or image. Thus, both tools do not take into account potentially more complex patterns. SarcApp (Neininger-Castro et al., 2023), Sarctrack (Toepfer et al., 2019), Sarc-Graph (Zhao et al., 2021), and ZlineDetection (Morris et al., 2020) give access to the length of individual sarcomeres. SarcApp uses deep learning to extract the features of complex patterns, which requires some programming skills and a computer compatible with deep learning. Sarctrack, Sarc-Graph, and ZlineDetection can analyze movies of moving patterns, but they cannot extract complex features. Sarctrack and Sarc-Graph do not offer a graphical user interface (GUI), and although ZlineDetection, has a GUI, it will require a minimum of coding knowledge to run it on the user’s computer. Except for SarcOptiM, all these tools do not require a manual selection from the user. Compared to these existing tools, PatternJ is very simple to use thanks to the GUI of ImageJ/Fiji and does not require programming skills. It can extract the position of complex features and the length of a single pattern. Additionally, it can combine the extraction results of multiple selections and display the distribution of spatial periods. Finally, it computes an image of the average pattern. Current PatternJ limitations are the requirement of user input for region selection and the incompatibility to analyze movies. The analysis of movies could be done currently by analyzing the same selection, one image at a time, a process that could be automatized in future versions of PatternJ. In conclusion, we believe our tool provides a useful ready-to-use solution to extract simple and complex features from biologically relevant patterns.

**Table 1.**
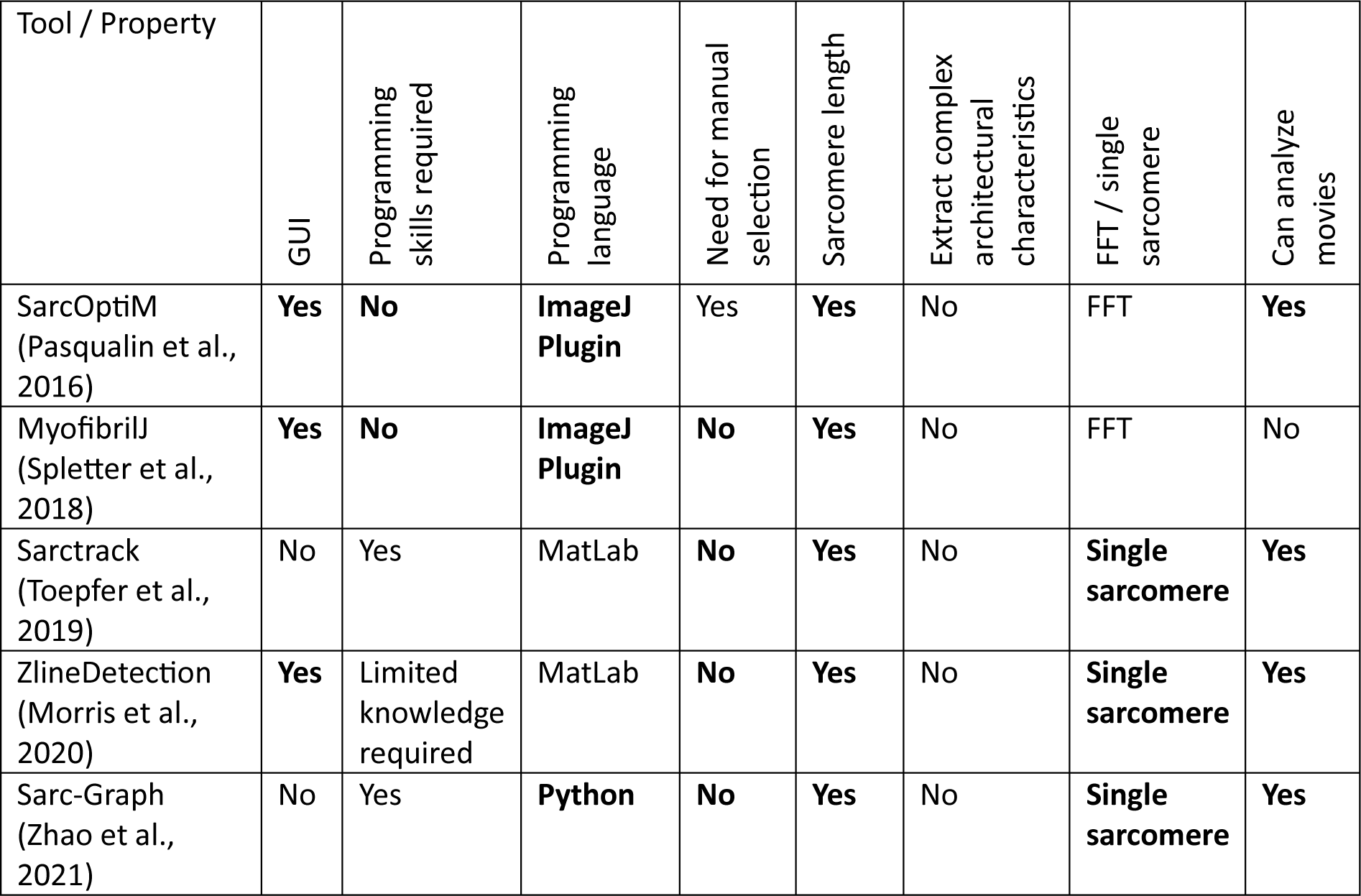
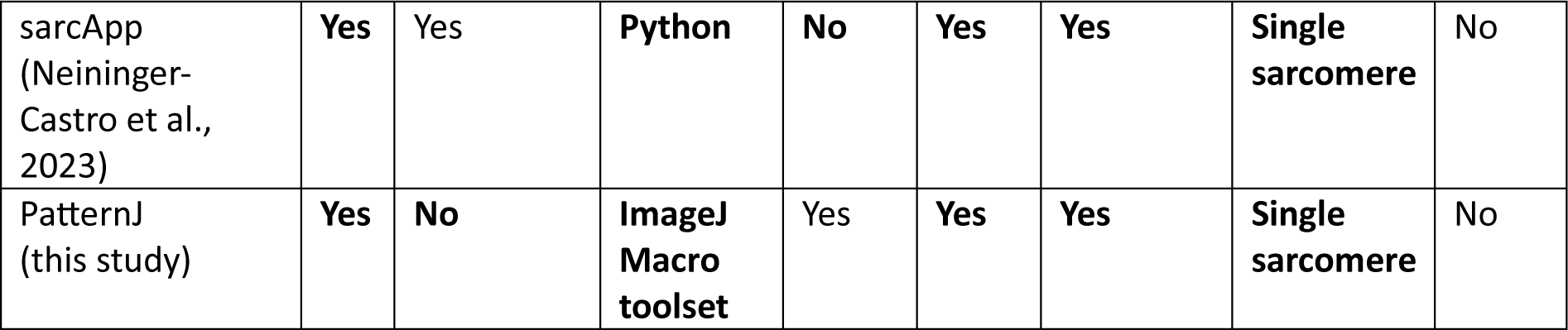

| Tool / Property | GUI | Programming skills required | Programming language | Need for manual selection | Sarcomere length | Extract complex architectural characteristics | FFT / single sarcomere | Can analyze movies |
| --- | --- | --- | --- | --- | --- | --- | --- | --- |
| SarcOptiM (Pasqualin et al., 2016) | Yes | No | ImageJ Plugin | Yes | Yes | No | FFT | Yes |
| MyofibrilJ (Spletter et al., 2018) | Yes | No | ImageJ Plugin | No | Yes | No | FFT | No |
| Sarctrack (Toepfer et al., 2019) | No | Yes | MatLab | No | Yes | No | Single sarcomere | Yes |
| ZlineDetection (Morris et al., 2020) | Yes | Limited knowledge required | MatLab | No | Yes | No | Single sarcomere | Yes |
| Sarc-Graph (Zhao et al., 2021) | No | Yes | Python | No | Yes | No | Single sarcomere | Yes |
| sarcApp<br>(Neininger-Castro et al., 2023) | Yes | Yes | Python | No | Yes | Yes | Single sarcomere | No |
| PatternJ<br>(this study) | Yes | No | ImageJ<br>Macro<br>toolset | Yes | Yes | Yes | Single sarcomere | No |

### Applications of PatternJ to biological samples

We validated our tool on simulated data, thus observing how it can analyze real data is important. To this end, we tested our tool with images acquired with various imaging methods in a wide range of tissues.

The main application of PatternJ will be to analyze images of muscle sarcomeres, from which extracting the position of proteins or protein domains is crucial to understanding how muscles are organized. We started by using our tool on confocal immunofluorescence images of cardiomyocytes in which sarcomeres were labeled for α-actinin (Taniguchi et al., 2009) (Fig. 4A). α-actinin is located at Z-disks displaying a single band (Figs. 4A and 4B). PatternJ can extract all positions of α-actinin on the selections chosen, from which it extracted the sarcomere lengths (Figs. 4B and 4C). Thanks to the subpixel capabilities of PatternJ, the distribution of sarcomere length shows a variety of contraction states within the cell. This first biological example shows that PatternJ can extract robustly a biologically generated single band pattern with high precision.

**Figure 4:**
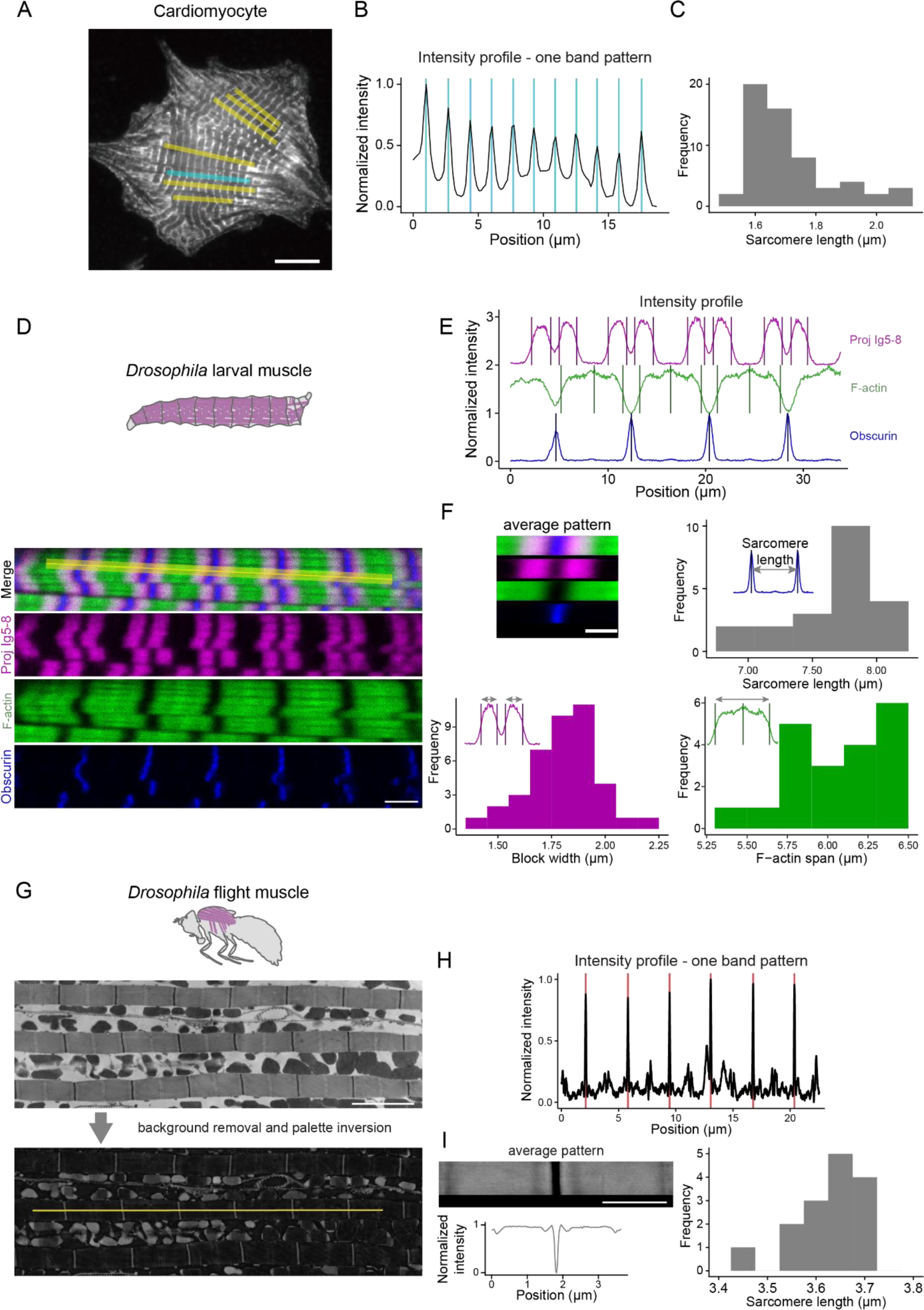
Application of PatternJ to images of muscle samples. (**A**) Confocal image of a cultured rat cardiomyocyte labeled for α-actinin adapted from (Taniguchi et al., 2009). (**B**) Intensity profiles obtained at the cyan selection in (A). Vertical bars correspond to the positions of bands extracted by PatternJ. (**C**) Distribution of sarcomere lengths from the yellow and cyan selections in (A). (**D**) Confocal image of a *Drosophila* larval body wall muscle with labeled Projectin (magenta, Proj-Ig5-8, Nano30 (Loreau et al., 2023)) and F-actin (phalloidin, green), and GFP-tagged Obscurin (blue). (**E**) Intensity profiles obtained at the yellow selection in (D). Vertical bars correspond to the positions of bands or blocks extracted by PatternJ. (**F**) Results of the analysis of 6 selections in the confocal image series of the larval muscle. Top left, image of the average pattern and average intensity profiles in each channel. Top right, distribution of the sarcomere length obtained from the position of Obscurin. Bottom left, distribution of the width of Projectin localization. Bottom right, distribution of thin filaments (F-actin) domain spanning. (**G**) Top, transmission electron microscopy image of adult *Drosophila* flight muscle (Loison et al., 2018). Bottom, same image background-corrected and with inverted pixel values. (**H**) Intensity profiles obtained at the yellow selection in (G). The red vertical lines correspond to the positions extracted by PatternJ. (**I**) Left, image of the average pattern and average intensity profile (in both cases the intensity levels were inverted to display like a traditional electron microscopy image). Right, distribution of sarcomere lengths from the 3 myofibrils in (G). Scale bars: (A) 10 µm, (D) and (G) 4 µm.

We continued with using a multicolor confocal image series of a *Drosophila* larval body wall muscle (Fig. 4D), displaying sarcomeres in which Obscurin-GFP is expressed and labeled for Projectin and F-actin (Loreau et al., 2023). These proteins display a variety of features: Projectin staining consists of two blocks, F-actin is a block with a bright band in its middle, Obscurin is a single band on the M-band (Fig. 4D). PatternJ extracts the edge of each of the block-like shapes and the position of the center of each band (Fig. 4E). Moreover, the tool provides an image of the average pattern, an average intensity profile, and the distribution of sarcomere length (Fig. 4F). From the positions extracted, we could additionally obtain the width of Projectin localizations and the length of the actin filaments (F-actin) (Fig. 4F). This example shows that PatternJ can reliably extract features or patterns that are more complex than a single band. The analysis in a multicolor image opens the possibility to observe how features in different channels vary at the pattern level.

Electron microscopy is another frequent means of imaging the organization of muscle. However, the images obtained are rarely quantified, because such images are challenging to analyze. We used an image of an adult *Drosophila* flight muscle (Loison et al., 2018) (Fig. 4G). These images display a very clear sarcomeric pattern, with the protein-rich Z-disk highlighted as a dark band. To extract the position of the Z-disk with PatternJ, one has to invert the image lookup table, so that it is treated as a bright band, similar to the previous examples using fluorescence microscopy. After the inversion and background correction, the positions of Z-disks are well extracted (Fig. 4H). PatternJ provides an average image of the pattern, an average intensity profile, and the distribution of sarcomere length (Fig. 4I). We show with this example that PatternJ is not limited to the analysis of images coming from fluorescence microscopy. More complex patterns from electron microscopy images may also be used with PatternJ.

Regularly spaced actin structures are not restricted to muscles. Actin rings were also observed in the axon initial segment (Leterrier et al., 2015; Xu et al., 2013). We tested PatternJ on super-resolved images of these actin rings obtained with the super-resolution method STORM (Fig. 5A) (Zhong et al., 2014). PatternJ extracts efficiently the position of individual actin rings. We find the typical 190 nm distance between actin rings that has been reported (Leterrier et al., 2015; Xu et al., 2013). Interestingly, we can also observe a significant variation around this value of about 28 nm (using the standard deviation, Fig. 5B). This variation of distance between rings can be noticed visually, potentially stemming from different protein organizations. This feature is rarely reported, because of a lack of tools to extract such data. Hence, we think our tool will be useful when analyzing such an organization; the access to the variability of distances between rings will be particularly useful when studying perturbations of the organization. The transformation of a table of protein localization obtained from STORM or other pointillist methods into pixelated images makes the use of PatternJ straightforward for this type of super-resolution imaging technique.

**Figure 5:**
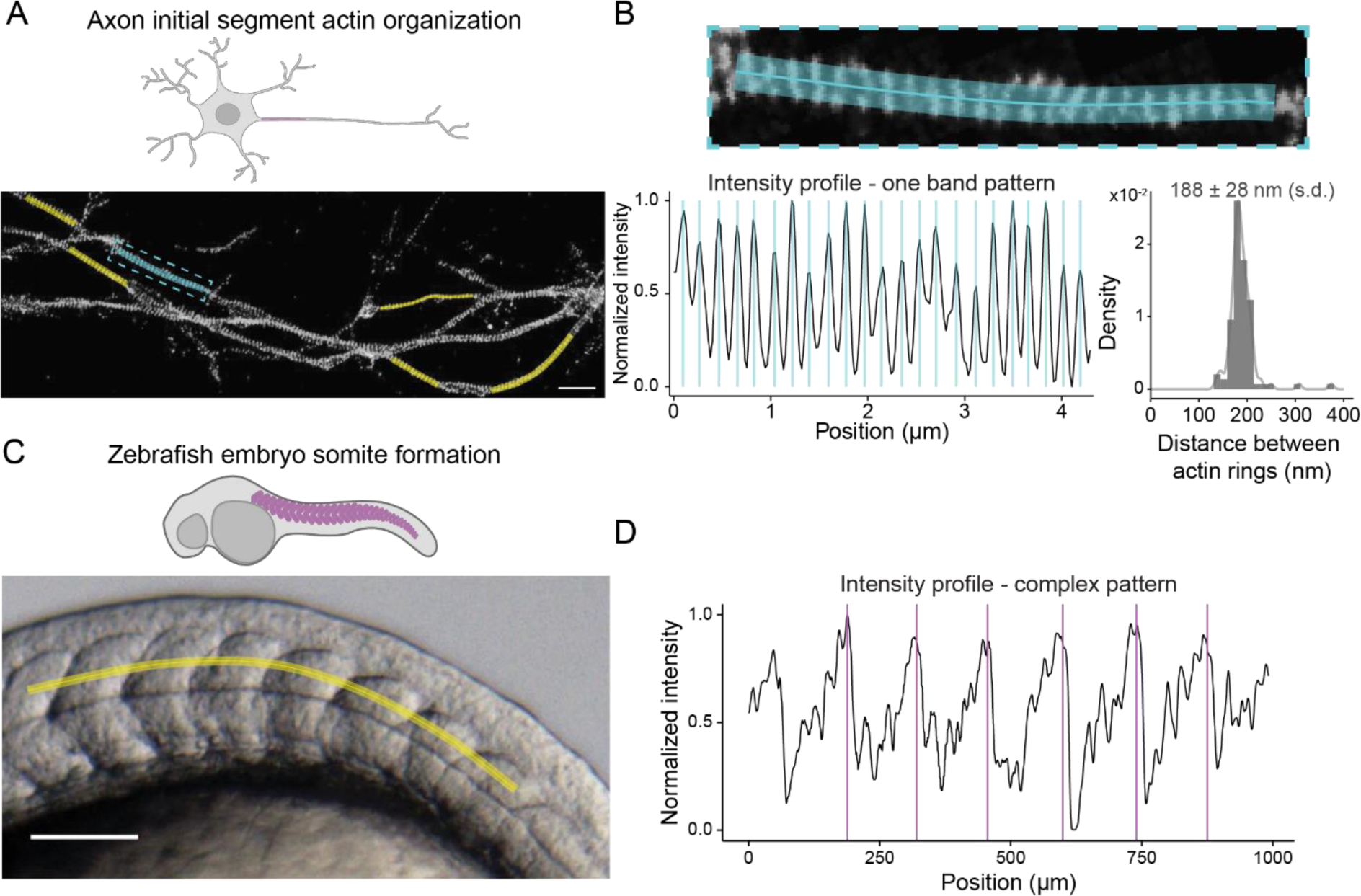
Application of PatternJ to images of biological samples. (**A**) STORM image of neurons, localizing actin filaments adapted from (Zhong et al., 2014). (**B**). Top, zoom in on the selection highlighted in cyan in (A). Bottom left, intensity profile obtained at the cyan selection in (A). Vertical bars correspond to the positions of bands extracted by PatternJ. Bottom right, distribution of sarcomere lengths obtained from the cyan and yellow selections in (A). (**C**) Bright-field image of a zebrafish embryo adapted from (Richter et al., 2017). (**D**) Intensity profile obtained at the yellow selection in (C). Vertical bars correspond to the positions of patterns extracted by PatternJ. Scale bars: (A) 2 µm, (C) 200 µm.

Patterns can take very complex shapes, which are difficult to describe with a simple mathematical function, limiting the potential extraction of features. Such patterns are however important to analyze. A bright field image of a zebrafish embryo is one example of such a situation: when extracting the intensity profile along formed somites, we obtained a very complex pattern, which is in part due to the presence of numerous granules (Fig. 5C). Our precise fitting algorithms for bands or blocks did not work in this condition. It is nevertheless possible to extract robustly the position of patterns (Fig. 5C), by using the second step of our algorithm, which identifies local maxima in the cross-correlation between the intensity profile and a single reference pattern profile. For this kind of situation, the extraction is not as precise as in the other examples, likely in the pixel range, but it is fully automated and it will be useful in situations in which many other algorithms would fail.

## Discussion

PatternJ is a simple-to-use toolset for ImageJ/Fiji that can extract complex features in images of repeated patterns. It can even extract features from challenging images containing high noise, for which it outperforms the precision of FFT or autocorrelation-only approaches to determine the average spatial period of the pattern (Fig. 2). The tool performs well in situations in which the intensity of the pattern varies or the variation of the spatial period does not exceed 20% (Fig. 3). The tool can extract geometric features of the sarcomeric pattern from muscle samples and other tissues displaying repeated patterns (Figs. 4 and 5). Even if it is designed primarily for fluorescence microscopy images, it can also be used on images taken with electron microscopy or bright field microscopy (Figs.4C and 5B).

Several tools to analyze such images already exist. What makes PatternJ particularly useful for the biological community? The main advantage of PatternJ compared to other tools is that it does not require any programming knowledge or any specific computer and yet, it can extract complex pattern features. In contrast to other tools that offer the automated selection of regions of interest, PatternJ does require the user to select the region of interest manually. However, regions that are selected automatically in other tools are likely to eventually require manual curation. Except for the manual selection, the other steps in PatternJ are automated. Compared to FFT or autocorrelation-only algorithms, above an SNR of 3, PatternJ is not advantageous if one is only interested in the average length of the spatial period. However, PatternJ provides the complete distribution of spatial period length, containing additional information to understand the variability in the pattern organization or even multiple modes in the length distribution that would otherwise be lost using only the average period length.

What are the limitations of PatternJ? Compared to some of the existing tools, PatternJ does not allow for the analysis of temporal sequences. Currently, one would have to use the tool sequentially on the images of the temporal sequence. This could be solved in a future version of the tool. Another limitation can appear in low SNR conditions when a pattern consists of multiple bands. Currently, PatternJ will search in a given pattern for the number of bands indicated by the user, by looking for local maxima in the intensity profile. However, in low SNR conditions, noise will generate local maxima that may be wrongly identified as bands. A simple solution to this limitation is to use Gaussian blurring, which is a built-in PatternJ function, or deconvolution for more advanced users: noise is then significantly reduced and multiple-band patterns are then usually well extracted. When possible, using a thicker linewidth in the selection can also mitigate such limitation as it reduces the effect of noise (Fig. 2B). Finally, electron microscopy images can be used, but since relevant information is usually in electron-dense regions, where pixel values are low, it will be beneficial for the user to invert the pixel values to make them analogous to fluorescence images. After this simple step, which is included in PatternJ functions, pattern features will be extracted with high precision.

Thanks to its ease of use and its capabilities, PatternJ is aimed at being used by a wide range of users, in fields that study the formation of regular biological patterns. Its precision and automation make analysis fast, robust, and reproducible, which we hope will help speed up discoveries.

## Methods

### *Drosophila* preparation for fluorescence microscopy

Larval muscles were stained with fluorescent nanobodies (Loreau et al., 2023) and rhodamine-phalloidin (1:1000 Molecular Probes) for 2 hr at RT or overnight at 4 °C as previously described in detail (Loreau et al., 2023). They were mounted in SlowFadeTM Gold Antifade (Thermo Fisher) and imaged with a Zeiss LSM880 confocal microscope using a 63× objective.

### *Drosophila* preparation for electron microscopy

The detailed protocol for how the electron microscopy of flight muscles was obtained was reported in (Loison et al., 2018). After staining and epon embedding, sections of 90 nm were cut on a Leica UC7 microtome and additionally stained with 2% UA for 30 min and 0.4% lead citrate for 3 min to enhance the contrast. Images were acquired with a Zeiss EM 900 (80 kV) using a side-mounted camera from Osis (Loison et al., 2018).

### Simulations

Simulated images used for the validation of PatternJ were generated in Python. In brief, single bands were added to oversampled images, that were then blurred with a Gaussian function to simulate diffraction. After resampling to the pixel size of a typical microscope using a 100X objective (42 nm), we added Poisson noise to simulate electronic noise.

Images of varying band intensity were obtained in the following way. A given band has one value for its intensity, which is homogeneous in a band. When varying the intensity, bands take values for their intensity in a normal distribution, of mean value 1 and of standard deviation that depends on the simulation, ranging from 0 (all bands have the intensity 1) to 0.25 (band intensity typically takes value in the 0.5 – 1.5 range). To vary band positions, each band could shift its position by selecting randomly a shift amplitude in a normal distribution of mean 0 and of increasingly large standard deviation. After the generation of positions, images were classified based on the difference in percent of the distances between the two bands closest and the two bands furthest.

A band was considered detected if the algorithm could find a band within 2.36 pixels from its actual position (corresponding to 100 nm at a 42 nm pixel size). The code used to generate these images is accessible at **<u>github.com/PierreMangeol/PatternJ</u>**.

## Availability

PatternJ can be downloaded from **<u>sites.google.com/view/patternj</u>**. The website also contains a tutorial to install and use the tool.

## Acknowledgments

We would like to acknowledge the members of the Schnorrer lab for discussion and feedback on PatternJ. This work was supported by the Centre National de la Recherche Scientifique (CNRS, F.S.) Aix-Marseille University (P.M.), the European Research Council under the European Union’s Horizon 2020 Programme (ERC-2019-SyG 856118 to F.S.), the France-BioImaging national research infrastructure (ANR-10-INBS-04-01) and by funding from France 2030, the French Government program managed by the French National Research Agency (ANR-16-CONV-0001) and from Excellence Initiative of Aix-Marseille University - A*MIDEX (Turing Centre for Living Systems).

