## Supporting information for "PatternJ: an ImageJ toolset for the automated and quantitative analysis of regular spatial patterns found in sarcomeres, axons, somites, and more"

This PDF file includes:

Figures S1 to S3

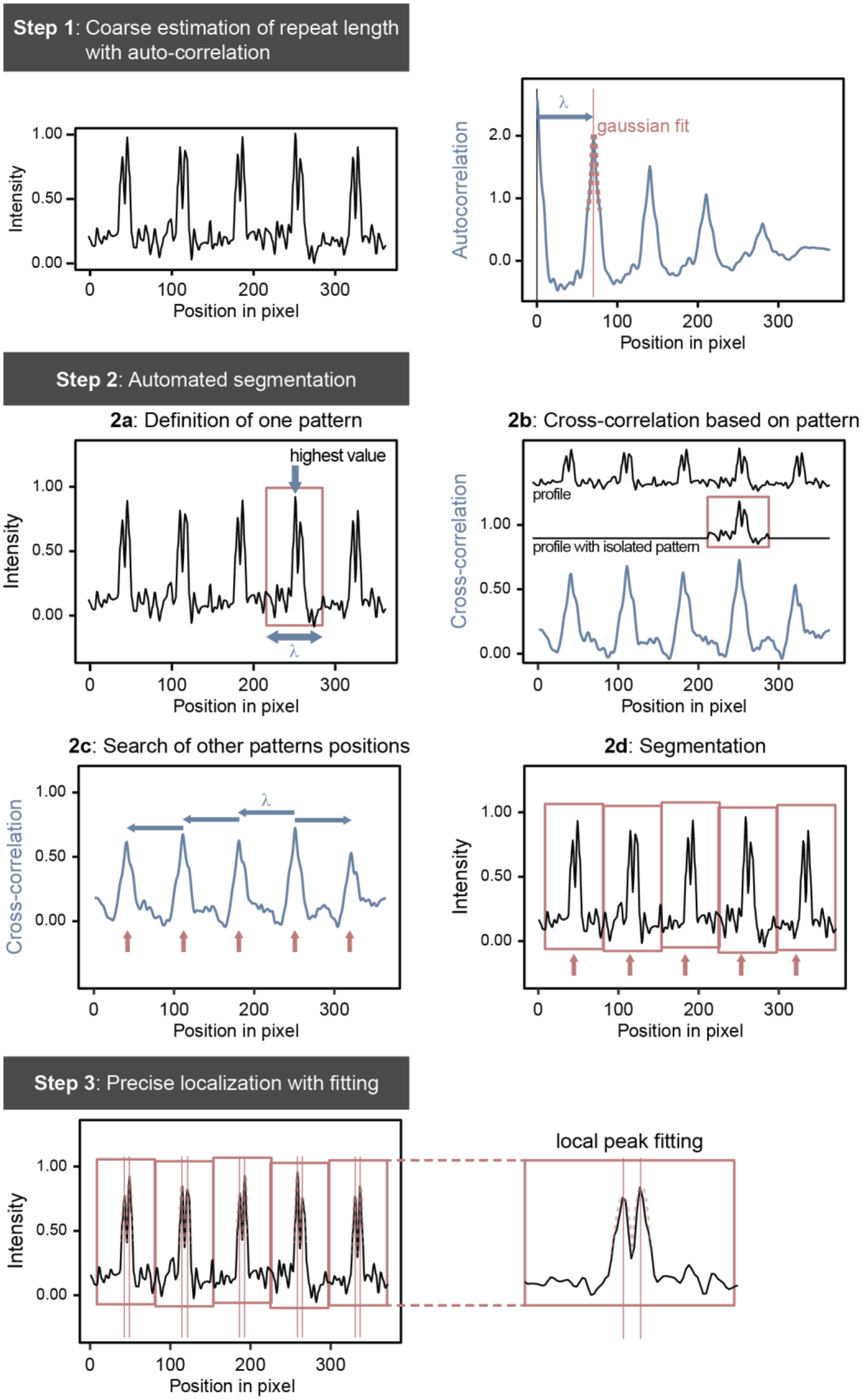
Step 1: Coarse estimation of repeat length with auto-correlation
Intensity
Position in pixel
λ
gaussian fit
Autocorrelation
Step 2: Automated segmentation
2a: Definition of one pattern
highest value
2b: Cross-correlation based on pattern
profile
profile with isolated pattern
Cross-correlation
2c: Search of other patterns positions
2d: Segmentation
Step 3: Precise localization with fitting
local peak fitting

**Figure S1**: Main steps followed by PatternJ to automatically extract pattern features precisely. Complete description in the main text.

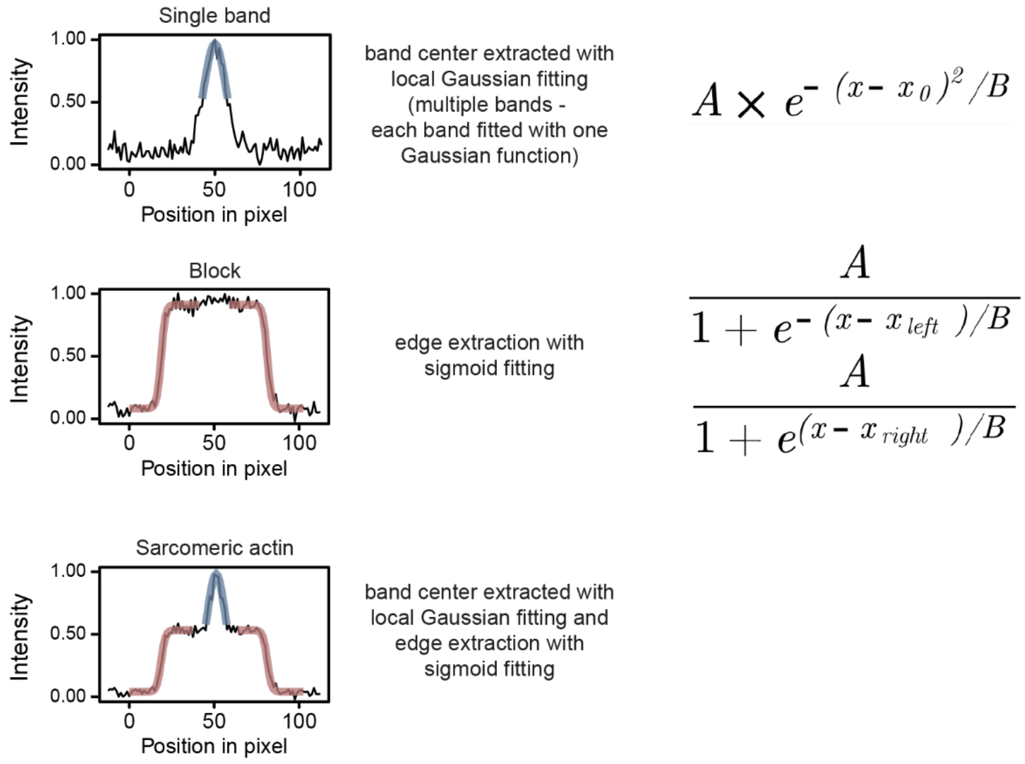


**Figure S2**: Fitting procedures used to extract pattern features. Features identified as “bands” are fitted with a Gaussian function. “Blocks” are fitted with two sigmoid functions. “Sarcomeric actin” pattern is fitted by one Gaussian function in its center and one sigmoid at each edge of the pattern.

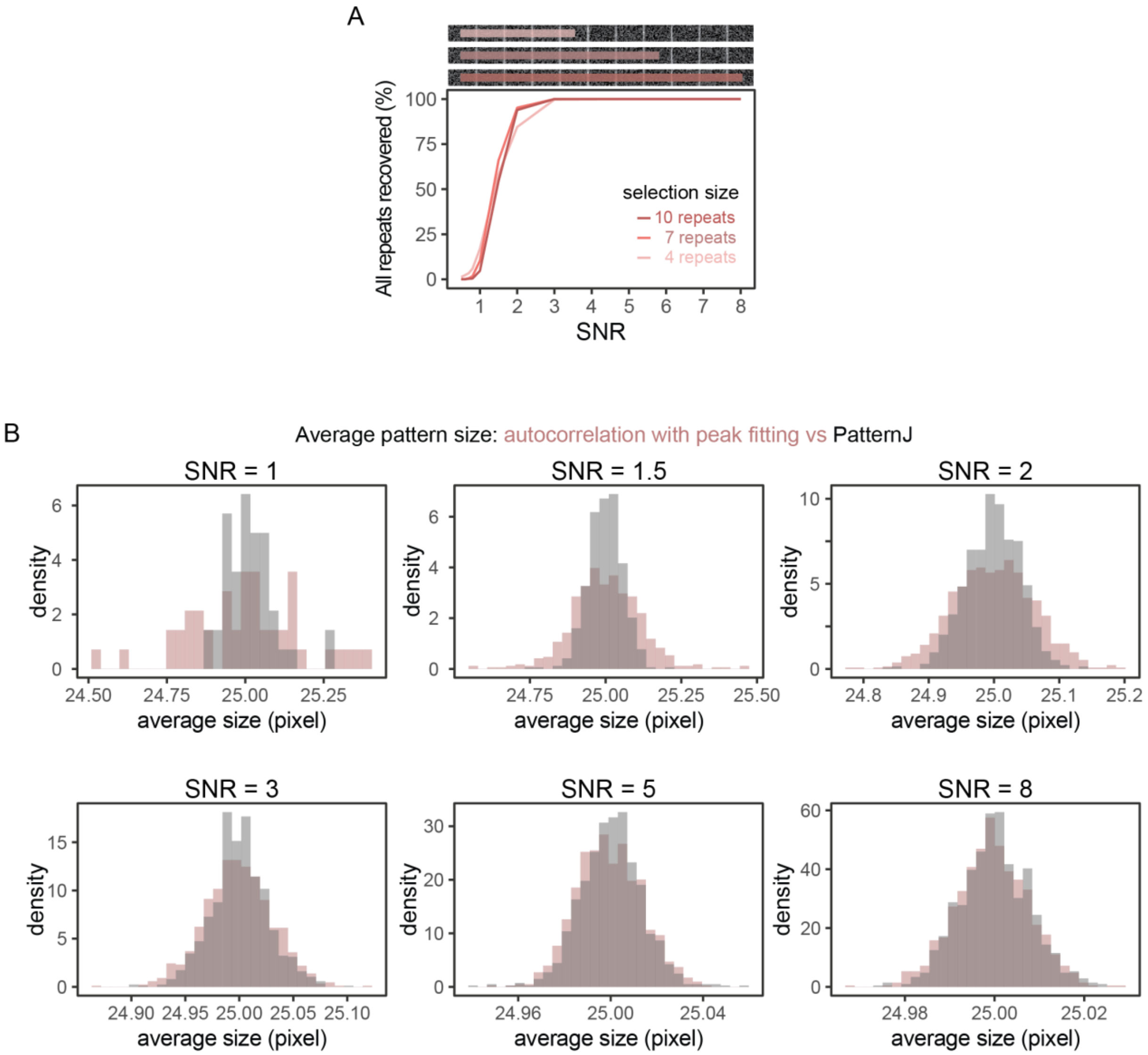


**Figure S3**. (A) Effect of the selection length on the performance of PatternJ to extract all patterns from a selection. (B) Comparison of the precision of PatternJ and autocorrelation using peak fitting, for a range of SNR typically observed on images of biological samples.
